# Compositional Data Analysis using Kernels in Mass Cytometry Data

**DOI:** 10.1101/2021.05.08.443265

**Authors:** Pratyaydipta Rudra, Ryan Baxter, Elena WY Hsieh, Debashis Ghosh

## Abstract

**Motivation:** Cell type abundance data arising from mass cytometry experiments are compositional in nature. Classical association tests do not apply to the compositional data due to their non-Euclidean nature. Existing methods for analysis of cell type abundance data suffer from several limitations for high-dimensional mass cytometry data, especially when the sample size is small.

**Results:** We proposed a new multivariate statistical learning methodology, Compositional Data Analysis using Kernels (CODAK), based on the kernel distance covariance (KDC) framework to test the association of the cell type compositions with important predictors (categorical or continuous) such as disease status. CODAK scales well for high-dimensional data and provides satisfactory performance for small sample sizes (*n* < 25). We conducted simulation studies to compare the performance of the method with existing methods of analyzing cell type abundance data from mass cytometry studies. The method is also applied to a high-dimensional dataset containing different subgroups of populations including Systemic Lupus Erythematosus (SLE) patients and healthy control subjects.

**Availability and Implementation:** CODAK is implemented using R. The codes and the data used in this manuscript are available on the web at http://github.com/GhoshLab/CODAK/.

**Supplementary information:** Supplementary Materials.pdf.

## 1 Introduction

Recent developments in single-cell based technologies, such as mass cytometry (i.e. cytometry by time-of-flight, CyTOF), has led to the need for computational and analytic approaches that can accommodate the high dimensionality and single-cell granularity. The analysis of CyTOF data can elucidate novel disease biomarkers and mechanisms of the underlying immunopathology, leading to improved treatments and prognostic measures.

Mass cytometry allows the simultaneous detection of more than 40 proteins per cell in hundreds of thousands of cells per sample (Saeys *et al*., 2016; Bendall *et al*., 2011). The data are often clustered into cell sub-populations first, which can then be used to answer scientific questions regarding the abundance of cell types and expressions of specific parameters (e.g. activation markers, signaling proteins, cytokines) across groups, such as disease and control groups, pre- and post treatment groups, samples that are stimulated or not. There have been significant research on clustering procedures with these high-dimensional data sets (see Aghaeepour *et al.* (2013) and Weber and Robinson (2016) for a review). We will focus on the downstream statistical analysis after the clustering has been performed. The statistical questions about the tree-structured cell population data (See Figure 1, for example) can be visualized in two layers. First, it is clinically interesting to know if the abundance of the cell subpopulations is different across two or more groups and/or conditions. Given the proportion of cell types for each sample, the next question is whether there is any differential expression of activation markers, signaling proteins, or cytokines (functional measurements of the cell populations studied). The latter is also known as ‘cell state’ analysis while the former is called ‘cell type’ analysis (Weber *et al*., 2019). In this paper, we focus on the analysis of cell type.

**Figure 1:**
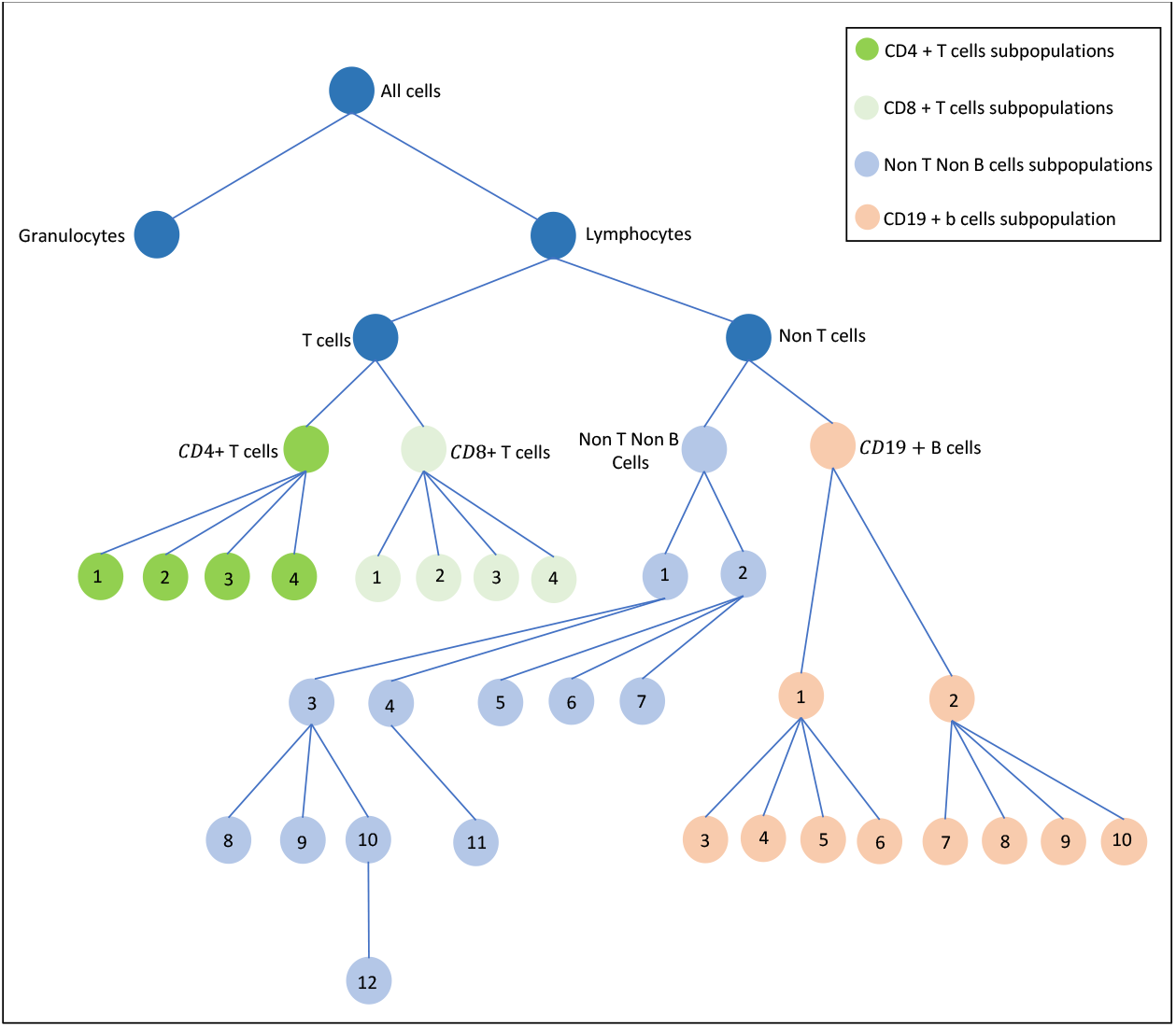
Hierarchical tree structure of the cell types from the SLE study 2. See Supplementary Materials for a full list of the cell subpopulations. It is of interest to (i) test if the compositional profile of the cell types is associated with the disease groups, i.e. if there is differential abundance of any of the cell types between SLE patients and controls; and (ii) if yes, which cell types contribute the most to the association.

While there is a variety of methods that test differential cell type abundance (Bruggner *et al*., 2014; Arvaniti and Claassen, 2017; Lun *et al*., 2017; Weber *et al*., 2019), several of them suffer from limitations such as

1. Difficulty of interpretation due to conflating cell type (abundance/frequency) and cell state (activation, function) questions.
2. No way of testing statistical significance for individual features (cell types).
3. Computational scalability.

See Weber *et al.* (2019) for a more detailed discussion on these limitations. The two approaches that overcome these limitations are (i) a GLMM-based approach using logistic mixed model (Nowicka *et al*., 2017) and (ii) the diffcyt method by Weber *et al.* (2019). These approaches effectively perform mass univariate analyses for each cell type. This ignores correlations between cell types as well as increases the multiple testing burden. In this paper, we propose a statistical framework that can test for the association of the multidimensional cell-type profile (or, ‘cell-type abundance’, to be used interchangeably) with the predictor variables (e.g. disease groups). Starting the analysis with the test of this global null hypothesis avoids the problem of multiple comparisons and also accounts for the correlation structure present in the data.

This multivariate approach can often help us better understand the biological functions of immune cell types. In the context of understanding disturbances to normal immune phenotype and function (i.e. a disease process, iatrogenic intervention), the interdependent relations between different cell types and their function need to be addressed collectively. If we evaluated statistical significance for changes in each single cell population, independent of the frequency changes in other cellular populations, it would not accurately depict the immunopathology, as the immune system is dynamic and changes in one compartment directly (or indirectly) affect the other. This is also the reason why single disease biomarkers do not accurately prognosticate disease progression or response to therapy, but “biomarker panels” are currently used, as single parameters cannot account for the underlying pathology (no matter how significant that single parameter is). Lastly, the field of multi-omic and systems immunology is accelerating our understanding of disease processes because of the capability to assess multiple processes simultaneously, depicting a closer view of the disease process, and hence statistical analyses must account for this multiplexed capability as opposed to addressing single differences.

### 1.1 Motivating data

**Study 1:** This example originates from a previous CyTOF study conducted by O’Gorman *et al.* (2017) to understand single-cell phenotypic and functional characterization of pediatric SLE patients and healthy controls. SLE patients with presentation prior to age 18 years old and age and gender-matched controls were recruited for the study. Peripheral blood was collected at initial diagnosis prior to the initiation of any treatment. Blood samples were fixed either immediately after collection (T0); or after incubation at 37C with a protein transport inhibitor cocktail for 6 h (T6). The final data set after all filtering steps contained 28 single-cell data files (7 patients and 7 controls, and 2 stimulation conditions, T0 and T6, per subject). Single cell data for every subject was manually gated to obtain the hierarchical clustered cell type data. It is of interest to test if the cell-type compositions are different (i) across the disease groups (ii) across the stimulation conditions.

**Study 2:** A second study comparing SLE patients and healthy controls is currently being conducted by the authors with a larger number of participants (See Supplementary Materials Section S2, for a partial analysis of this data using our method). The hierarchical clustering structure for this study is shown in Figure 1. Due to the hierarchical nature of the data, cell-type abundance can be defined in many different ways by choosing different parent and children nodes of the tree. For example, it might be of interest to consider the abundance of all cell types from the terminal nodes as a fraction of the lymphocytes, but one may also want to test, for example, the abundance of the B-cell subpopulations as a fraction of the B-cells. Our proposed multivariate approach can be used to conduct the test to answer each of these questions separately.

### 1.2 Statistical challenge

Data on cell-type abundance is compositional by nature, i.e. the sum of the cell-type proportions add up to one. In mathematical notation, if *P ≡* (*P*_1_, *P*_2_, …, *P*_*q*_) denotes the cell-type abundance of the *q* cell types, then

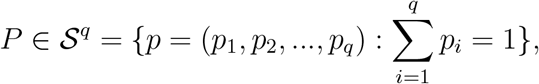

where *𝒮*^*q*^ is the *q*-dimensional simplex (Aitchison, 1982). Due to this, the classical statistical models for non-compositional data are not appropriate for testing differential abundance in mass cytometry.

Although there is a rich literature of statistical methods for compositional data analysis (Pawlowsky-Glahn *et al*., 2015), most of the traditional methods of compositional data analysis (often based on multinomial or Dirichlet distributions) test the association of the predictors with the individual components one at a time. A multivariate approach accounting for the correlation among the components is expected to perform better and have higher statistical power due to a decreased burden of multiple testing. The advantage of using such multivariate approaches for genomic association studies is well documented (e.g. Wu *et al*. (2011); Broadaway *et al.* (2016)). The authors are unaware of an appropriate multivariate approach to test association in high-dimensional cell-type abundance data arising from mass cytometry.

The traditional generalized linear models cannot be used here due to the presence of overdispersion typical for these data. The state of the art methods to analyze differential abundance for mass cytometry data are based on classical generalized linear mixed models (GLMM) (Nowicka *et al*., 2017) with the help of ‘observation level random effects’ (OLRE), or based on newer developments such as edgeR, limma or voom (Robinson *et al*., 2010; Ritchie *et al*., 2015; Law *et al*., 2014). The recently developed diffcyt methods (Weber *et al*., 2019) using the above approaches are shown to perform well for mass cytometry data, but they have the same limitation of approaching the problem in an univariate manner. Also, it has been shown that the statistical tests based on the GLMM approach often leads to anticonservative results, especially in small samples (Silk *et al*., 2020; Bolker *et al*., 2009; Forstmeier *et al*., 2017) which is typical for clinical studies using CyTOF data. The newer methods such as edgeR or voom do not provide theoretical guarantee of type-I error control either. These methods have been shown to have inflated type-I error rate when applied to other types of data (Hawinkel *et al*., 2019; Rocke *et al*., 2015; Vestal *et al*., 2020; Datta and Nettleton, 2014). In this paper, we propose a multivariate statistical framework, ‘Compositional Data Analysis using Kernels’ (CODAK), based on kernel distance covariance (KDC) (Hua and Ghosh, 2015) to quantify and test the association of predictors such as grouping or application of drugs with the composition profile of cell types. This association test can often be used as the test of differential abundance, but we present the method in general so that it can be used for more general cases (e.g. continuous predictor) besides the two-group comparison. We also propose two approaches of covariate adjustment under this framework and suggest some follow-up methods to understand which components of the composition are most responsible for the association. We illustrate the performance of our methods using extensive simulation studies and also apply it to high-dimensional mass cytometry dataset we collected on Systemic Lupus Erythematosus (SLE) patients and healthy control subjects. Analysis of the data revealed clinically relevant patterns such as differential cell type abundance between the disease and the control group.

## 2 Methods

### 2.1 The Kernel Distance Covariance framework

Distance covariance/correlation is a method to quantify and test for association between random variables of arbitrary dimensions (Székely *et al*., 2007, 2009). It is powerful against any form of lack of independence. The distance covariance approach is closely related to the kernel-based approaches using Hilbert Schmidt Independence Criterion (HSIC) (Gretton *et al*., 2007). The equivalence of the two approaches were shown by Sejdinovic *et al.* (2013) and Shen and Vogelstein (2020). Hua and Ghosh (2015) discussed the equivalence in the context of genetic association studies and introduced the term ‘kernel distance covariance’ (KDC).

For *n* measurements on two multidimensional random variables *X*_1*×p*_ and *Y*_1*×q*_, let us denote the observation from *i*th sample unit as (*X*^(*i*)^, *Y*^(*i*)^). Define the matrices *K* = (*k*_*ij*_) and *L* = (*l*_*ij*_) as

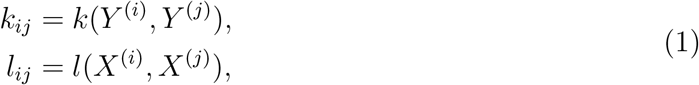

where *k* and *l* are the appropriate kernel functions measuring the similarity of pairs of observations. The KDC statistic is defined as

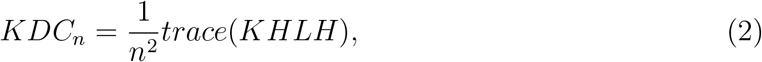

where 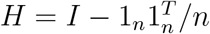 is the centering matrix, *I* being the identity matrix of dimension *n*, and 1_*n*_ being the *n* × 1 vector with each element equal to 1.

For our application of the KDC approach, suppose we have a (potentially multivariate) predictor *X* = (*X*_1_, *X*_2_, …, *X*_*p*_) and that the cell-type abundance is given by *P* = (*P*_1_, *P*_2_, …, *P*_*q*_), where 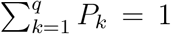. A key aspect of the KDC approach is the choice of the kernels *k* and *l*. Some common choices of kernels in association studies are the linear kernel, polynomial kernel, and Gaussian kernel (Schölkopf *et al*., 2004). However, these do not directly apply to cell-type abundance compositional data since the data belongs to a simplex. For CODAK, we propose a Gaussian kernel (Schölkopf *et al*., 2004) with Aitchison distance (Aitchison, 1982) as an appropriate kernel for measuring similarity between two compositions. Let the cell-type composition for the *i*th sample be *P*^(*i*)^ = (*P*_*i*1_, *P*_*i*2_, …, *P*_*iq*_). Then the similarity between the composition profiles of the *i*th and the *j*th sample can be given by:

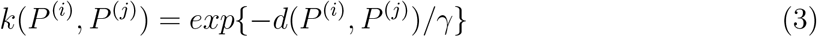

where *d*(*P*^(*i*)^, *P*^(*j*)^) is the Aitchison distance (AD) between *P*^(*i*)^ and *P*^(*i*)^, and *γ* is a tuning parameter which is often chosen as the median distance. The AD is defined by

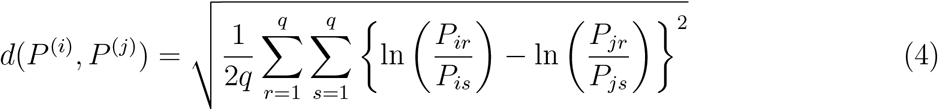

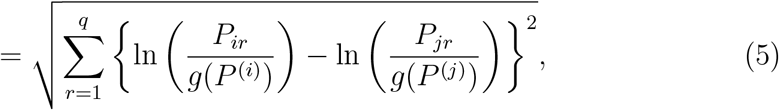

where *g*(*P*) is the geometric mean of *P*_1_, *P*_2_, …, *P*_*q*_. The AD has some desirable properties such as scale invariance, permutation invariance, perturbation invariance, and subcompositional dominance (See Martín-Fernández *et al.* (1998) for a discussion).

For most applications, *X*^(*i*)^ is univariate and for the remainder of the paper we treat it as such. We use a linear kernel for continuous predictors defined by

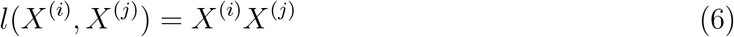

and a kernel based on Hamming distance for binary predictors given by

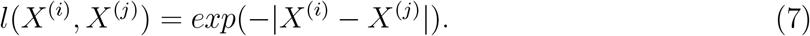

Since the Hamming distance is a distance metric, we can use standard arguments (Sejdinovic *et al*., 2013; Shen and Vogelstein, 2020) to show that (7) is a proper kernel. We use a permutation method to obtain the null distribution of the KDC statistic. We also explored the performance of a Gaussian kernel for *X* and found it to have slightly worse statistical power (results not shown).

The intuition for the KDC approach for the motivating data (study 1) is demonstrated by Figure 2. It is often the case that the cell type compositions are different in two disease groups such as SLE and healthy control. Loosely speaking, this is same as saying that the similarity in the compositional profiles between observations from the same disease group are likely to be more compared to that between observations from different disease groups. This is reflected in the first panel where the red ‘same group’ curve is located to the right of the blue ‘different group’ curve. On the other hand, the compositions are less likely to vary across stimulation conditions, which is why the curves in the second panel are more similar.

**Figure 2:**
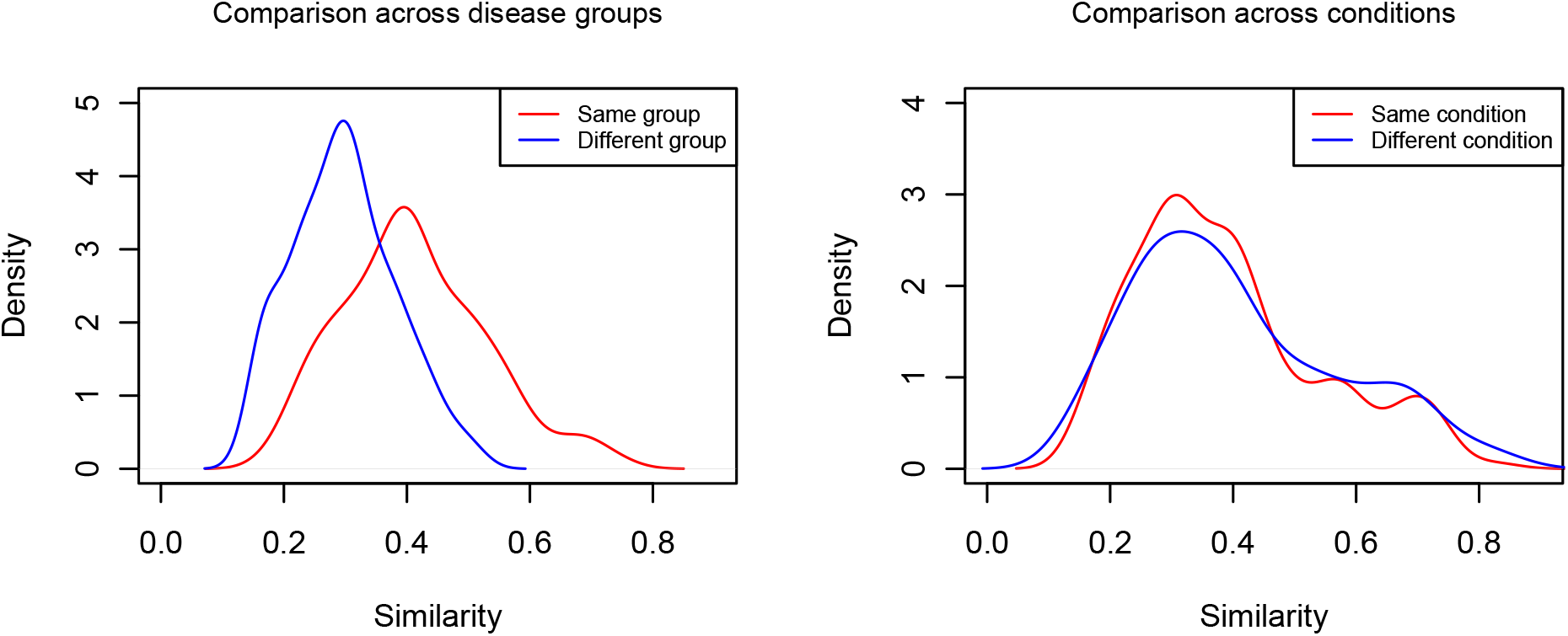
Motivation of the KDC approach for the first SLE study: The densities for the kernel similarity measures are plotted when comparing the two disease groups (first panel) and when comparing the two stimulation conditions. The red curve shows the similarity of observations within the same group (conditions) and the blue curve shows the similarity of observations between groups (conditions).

### 2.2 Adjusting for covariates

Our framework CODAK allows for covariate adjustment using two different approaches. Suppose we have a set of covariates *Z* = (*Z*_1_, *Z*_2_, …, *Z*_*K*_). Denote the value of *Z* from the *i*th subject as *Z*^(*i*)^. In the first approach, we use the additive log-ratio transformation (alr) commonly used in compositional data analysis (Aitchison, 1982; Pawlowsky-Glahn *et al*., 2015; Aitchison *et al*., 2000) to transform the data to *Y* = *alr*(*P*) *∈* ℝ^*q*−1^, perform covariate adjustment to compute the residuals *Y* (*Z*) using linear regression, and transform the residuals back to the simplex 𝕊^*q*^ using the inverse alr transformation: *P*(*Z*) = *alr*^−1^(*Y* (*Z*)). We can then use CODAK on *P*(*Z*) and *X*. The alr-transformation and the inverse alr-transformation are given by

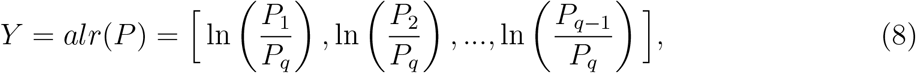

and

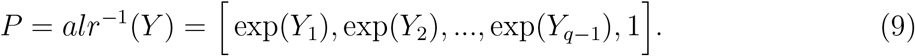

Here we can skip the technicality of converting *P* = *alr*^−1^(*Y*) to a composition such that 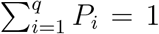 due to the scale invariance property of the AD. Note, however, that the alr-transformation is asymmetric in the components, and the alr-coordinates are non-isometric in nature (Pawlowsky-Glahn and Buccianti, 2011). Alternatively, one may use the centered log-ratio (clr) transformation and its inverse given by

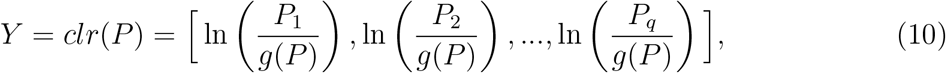

and

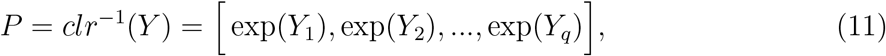

where *g*(*P*) is the geometric mean of *P*_1_, *P*_2_, …, *P*_*q*_. The clr transformed vector *Y* is symmetrical in the components, but belongs to a subset of ℝ^*q*^ due to the constraint 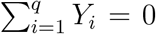. One can obtain an orthonormal coordinate system based on the clr-transformed vector, but it is not necessary for our purpose, and our empirical studies demonstrate that alr and clr have near identical performance for covariate adjustment. We present the results using the alr-based approach (CODAK-alr) in the Results section.

A second approach, motivated by Zhan *et al.* (2015), is developed when the covariates are categorical. This approach is based on the idea of stratified kernel, which, in our context, is defined as

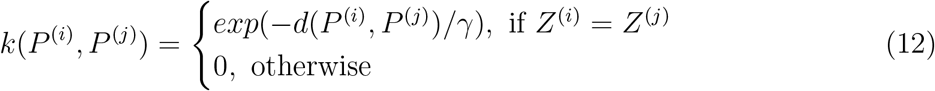

The kernel *l* is defined similarly. The method essentially considers two observations to have (potentially) positive similarity only if they are from the same stratum, where the strata are defined by the values of the categorical covariates. We can then proceed to use CODAK using Equation 2. It can be shown that the stratified kernel defined in this manner is strictly positive definite (Zhan *et al*., 2015; Park *et al*., 2012). It should be noted that this method reduces the effective sample size by only considering similarity within strata and therefore may lose power, especially in the presence of multiple categorical covariates. However, in many applications (e.g. the motivating problem), we only have one categorical covariate. The stratified kernel approach (CODAK-sk) can be effective in such cases, and has been empirically shown to be robust against slight violation of the independence of sample observations (See Section 3.3).

### 2.3 Follow up methods for individual components

One criticism of multivariate approaches is the difficulty of interpretation for individual components of the composition profile. For example, as a follow up to differential abundant cell-type compositions, it is often of interest to understand the cell types that are the top contributors to the differential abundance. While one can use the traditional effect sizes such as odds ratio or average difference in proportions for each cell type, we provide two kernel based solutions here.

#### 2.3.1 Leave-one-out approach

One intuition is that if a component *c* of the composition contributes to the association of the compositional profile *P* with a predictor *X*, then dropping that component from the compositional profile should lead to a decrease in the distance correlation. Therefore, we can compute the following leave-one-out (LOO) statistic for every component *c* ∈ {1, 2, …, *q*} and rank them in order of their values:

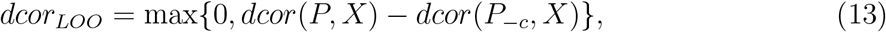

where *P*_−*c*_ = *P* (1, 2, …, *c* − 1, *c* + 1, …, *q*). It is obvious from 4 that the AD between two compositions *P*^(*i*)^ and *P*^(*j*)^ reduces when a component is excluded (subcompositional dominance). Further, it can be shown (see Supplementary Materials Section S3 for a simple proof) that the reduction is maximum when the component *c* with the 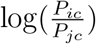 value farthest from the mean is excluded, i.e.

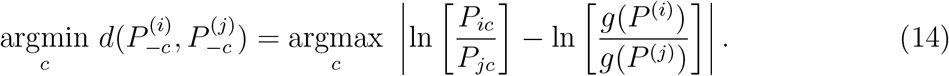

This further strengthens the above intuition since it is clear that the component with the highest true ratio of abundance in the two groups contributes the most to the AD between the composition profiles of observations in different groups. Therefore, we propose picking the top cell-type candidates based on the ranking of this LOO statistic defined in Equation 13.

#### 2.3.2 Weighted distance correlation approach

A second approach is motivated by Wen *et al.* (2020) and based on weighted Aitchison Distance (Egozcue and Pawlowsky-Glahn, 2016). Following Wen *et al.* (2020), define weighted distance correlation between *P* and *X* by modifying the kernel *k* in Equation 3 to use the weighted AD *d*_*w*_(*P*^(*i*)^, *P*^(*j*)^) instead of *d*(*P*^(*i*)^, *P*^(*j*)^), where the weighted AD is defined as

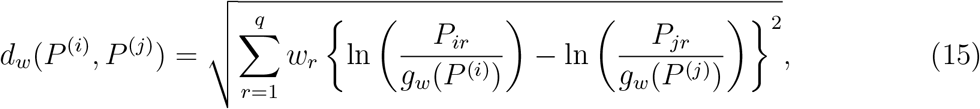

where 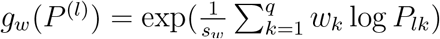 and 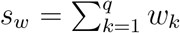. A set of weights that maximize the weighted distance correlation between *P* and *X* can then be obtained. Following Wen *et al*. (2020), we simplified the optimization problem by defining

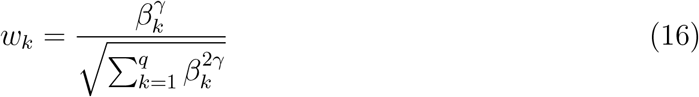

and optimizing for *γ >* 0, where *β*_*k*_ is the distance correlation between *P*_*k*_ and *X*. The larger the value of *γ*, it makes the weights of the most associated components contribute more to the weighted AD and subsequently to the KDC statistic. These weights can then be ranked in order of magnitude to indicate the top cell-type candidates contributing to the association.

### 2.4 Implementation

CODAK is implemented using statistical software package R, and the codes are available at http://github.com/GhoshLab/CODAK/. The time taken on a single computer to run CODAK for a mass cytometry data of usual size ranges from less than a second to approximately 90 seconds depending on number of permutations used for the test (10^3^ to 10^6^).

## 3 Results

### 3.1 Simulation studies

#### 3.1.1 Description of the simulations

We conducted simulations to compare type-I error and power performance of the different methods for the following scenarios: (i) no covariate (binary or continuous predictor) (ii) with covariate adjustment (binary or continuous predictor) (iii) with covariate adjustment for repeated measures (binary predictor). We report the results for the binary predictor (i.e. two group comparisons) here and the results for continuous predictor in the Supplementary Materials (Figures S1 and S3). The third scenario was explored to understand the robustness of the methods for the violation of the independence assumption.

For multivariate parameters, the effect size is hard to show in the power plots. While measures such as maximum log odds ratio (OR) or the true Aitchison distance can be used as effect size, we decided to explore the effect of them separately in Figure 4. Instead, for each of the above scenarios, we considered four cases: (a) no association (null hypothesis) (b) all cell types have some small association with the predictor (e.g. small difference in abundance of every cell type across two groups), (c) 50% of the cell types have small associations with the predictor (d) 25% of the cell types have larger associations with the predictor. The ‘small association’ is defined as the 0.2 percentage point difference, and ‘larger difference’ as 0.4 percentage point difference between the two group. For scenario (ii), the effect of the covariate *Z* is simulated similarly to the effect of the predictor in case (b). We used *N* = 10, 000 simulations for each case and plotted the estimated size and power against different choices of the nominal level *α*.

**Figure 3:**
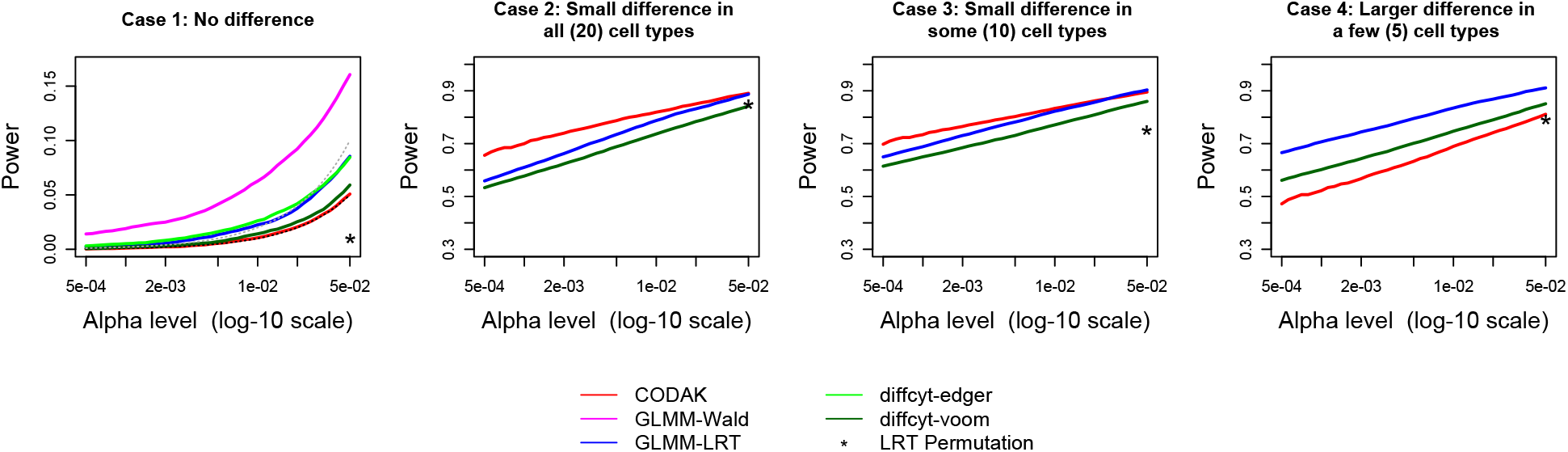
Comparison of statistical power for binary predictor. The black dashed line in the first plot shows the nominal level *α* and the grey dashed line shows two times *α*. Only the methods with reasonable control of type-I error are shown in the other 3 plots. The power for the LRT-permutation is shown for one choice of *α* due to the high computation time.

**Figure 4:**
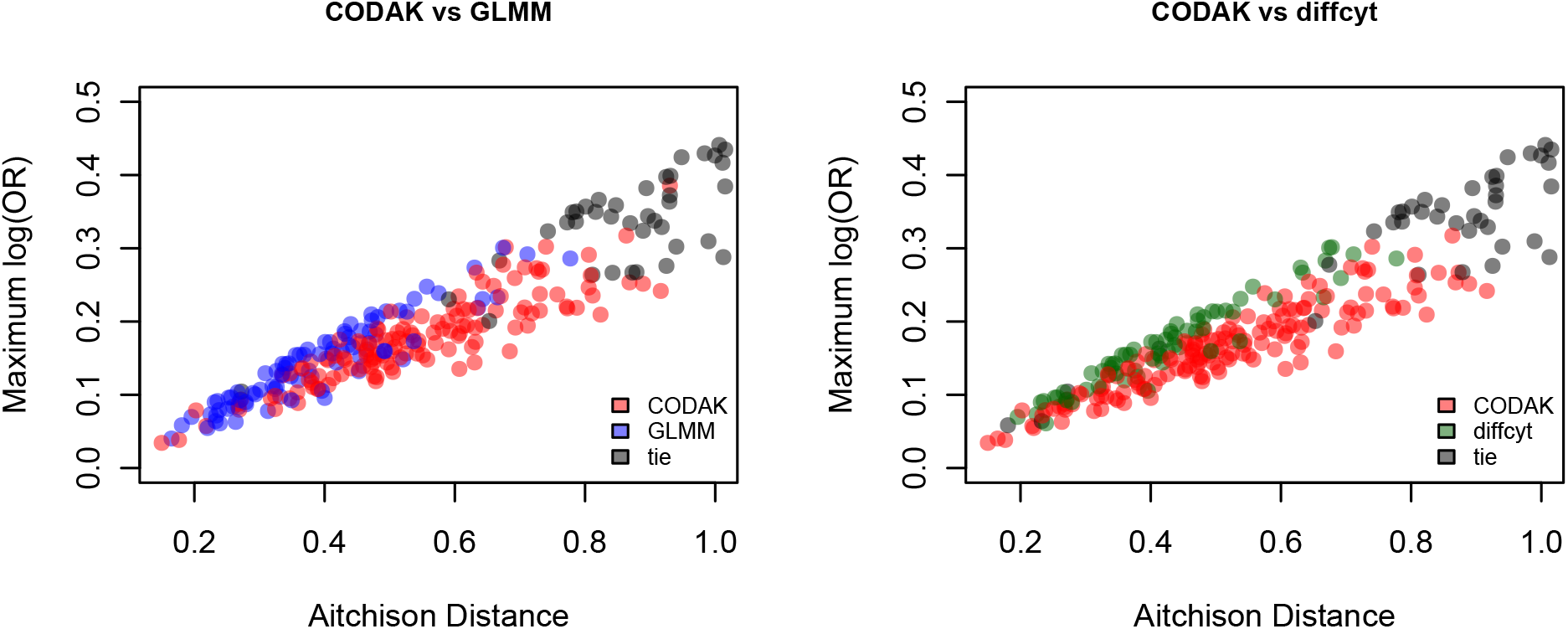
Comparison of CODAK with GLMM and diffcyt-voom for various effect sizes. For every simulation scenario, the true Aitchison distance (AD) and true maximum log odds ratio are plotted. The colors represent the method with a higher statistical power for that scenario. It is evident that CODAK favors higher AD while the other methods favors strong effects for individual components. Scenarios *AD >* 1 or | log(*OR*)| *>* 0.5 are not shown since all methods had perfect power in such cases.

We simulated data for 2*n* = 24 subjects, i.e. *n* = 12 per group for the two group comparison. Using a multinomial logit model, we generated 30, 000-50, 000 cells per individual sample with probability depending on the value of the predictor for a cell to be one of the 20 cell types. An observation-level random effects (OLRE) term was included in the multinomial logit model to model the overdispersion present in real CyTOF data sets. The parameters, such as probabilities in the control group (ranging from 0.002 to 0.15), and the standard deviation of the random effects term (*σ*_*b*_ = 0.2) were chosen based on the estimates obtained from real data (study 1). For scenario (iii), we used an additional subject-specific random effect term to induce the correlation in the repeated measures structure.

### 3.2 Competing methods

Besides CODAK, we considered two GLMM-based methods and two methods from the diffcyt package. The GLMM-based methods were both logistic mixed effects models to test the association of the predictor with the abundance of each cell type (followed by adjustments for multiple testing using Bonferroni method), the difference among the two methods being the test used: Wald test or LRT. The methods for diffcyt were also conducted per cell type (followed by adjustments for multiple testing) using either the edgeR or the voom method (as suggested by Weber *et al.* (2019)). It should be noted that there is also a GLMM test option in the diffcyt package which is essentially same as the Wald test we considered. Since both the GLMM-based methods resulted in inflated type-I error, we also applied a resampling method using the LRT statistic where the LRT statistic was calculated using the logistic mixed model and a permutation method was used to obtain the p-value instead of using the asymptotic distribution of the LRT statistic. Due to high computational time, we only used this method for one of our simulation scenarios and we only reported the power/type-I error rate for *α* = 0.05.

When adjusting for covariates, we included them directly in the model for the GLMM or diffcyt methods. For scenario (iii) with repeated measures, when fitting the GLMM models, we used the known information on which observations are coming from the same individual subject to fit multilevel models accounting for the repeated measures structure. On the other hand, CODAK did not use this information.

### 3.3 Results from the simulation studies

The results for binary predictor (Figure 3) shows that CODAK controls the type-I error at the target level, but none of the other methods could do so. The type-I error control of diffcyt-voom is near satisfactory and the LRT method among the GLMM methods leads to less inflation of the type-I error. Therefore, we choose one from each of the approaches of the competing methods and only show the diffcyt-voom and GLMM-LRT for the subsequent power comparisons. However, the comparison of power of GLMM-LRT and the other two methods are still unfair since the size of GLMM-LRT is approximately twice the as large as the target *α* level for the range of values of *α* considered. The power comparison shows that CODAK performed better than the other two methods in the case with small differences in all cell types (case b). With the reduction in the number of associated cell types, the relative advantage of CODAK compared to the univariate methods gradually diminished. The resampling-based LRT method controlled the type-I error at the nominal level, but failed to achieve high power. It is also extremely computationally intensive. Similar results were obtained for continuous predictor (See Suppementary Materials Figure S1).

The results for binary predictor in the presence of a covariate (Figure 5) lead to similar findings. CODAK-alr had slightly inflated type-I error here, but it was comparable to diffcyt-voom. CODAK-sk achieved good control of type-I error. However, CODAK-sk had inferior power performance compared to CODAK-alr. This is not unexpected, as discussed in Section 2.2.

**Figure 5:**
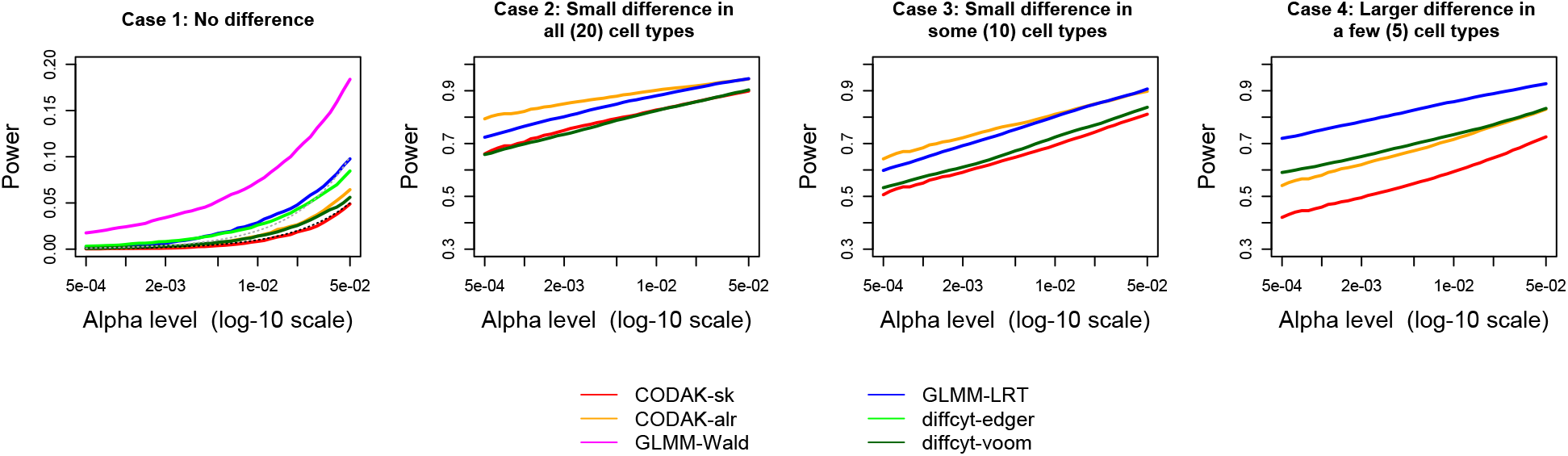
Comparison of statistical power for binary predictor adjusting for a binary covariate. The black dashed line in the first plot shows the nominal level *α* and the grey dashed line shows two times *α*. Only the methods with reasonable control of type-I error are shown in the other 3 plots.

The simulations from scenario (iii) are shown in Figure 6. CODAK-sk achieved reasonable control of type-I error under slight misspecification of of the independence assumption (subject-specific random effect variance = 0.1) and the power was also comparable to its performance in other cases. None of the other methods (including CODAK-alr) controlled the type-I error rate. We also verified that when the amount of dependence was increased, the control of type-I error by CODAK-sk gradually worsened (result not shown).

**Figure 6:**
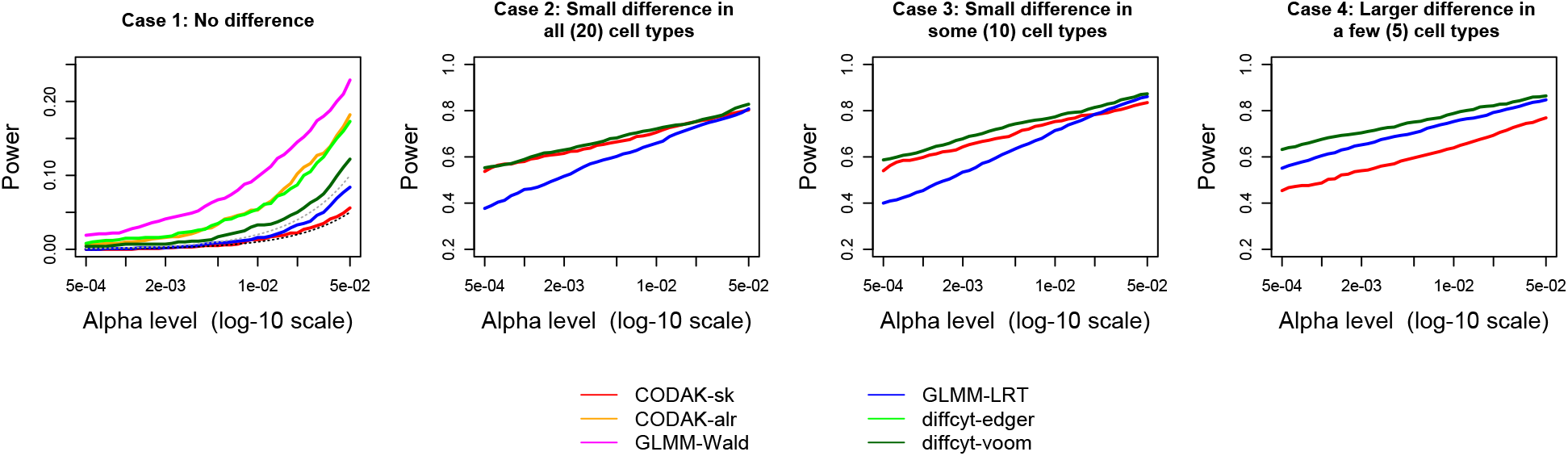
Comparison of statistical power for binary predictor adjusting for a binary covariate with repeated measures. The black dashed line in the first plot shows the nominal level *α* and the grey dashed line shows two times *α*. Only the methods with reasonable control of type-I error are shown in the other 3 plots.

To better understand the situations where CODAK (or other methods) have advantage, we plotted the true AD and maximum log(OR) for each simulation (from scenario (i)) and color coded according to which method had higher power (Figure 4). It is not unexpected that the GLMM and diffcyt-voom appeared to be perform better than CODAK when maximum log(OR) is high but the AD is not (blue/green points). These are the cases where a smaller number of cell types are associated and the association is strong. CODAK performed better for effects that are relatively weaker, but more spread out across the cell-types (red points). All methods performed well (black points) when there was strong association for several cell types, or very strong association for fewer cell types.

Finally, we compared the ranking of the cell types for the different follow-up methods. For every simulation, we calculated the rank correlation of the true log(OR) with the rank of the cell types obtained using the LOO and the weighted dcor methods. The median of the rank correlations was 0.52 for the LOO method and 0.41 for the weighted dcor method. These are reasonably high considering that the rank correlation between estimated log(OR) and the true log(OR) was 0.42. The median number of ‘true’ top-5 cell types in the top-5 list provided by these methods was 3 for all the three methods (LOO, weighted dcor, and estimated log(OR)). Based on these results, we can conclude that the performance of the LOO-based follow-up method to rank the contribution of the individual cell types is satisfactory and should be preferred over the weighted dcor method.

### 3.4 Real data

In order to study the performance of CODAK on real data we applied it on the SLE data from study 1 in Section 1.1. SLE is a systemic inflammatory disease in which multiple immunological derrangements have been described – Toll-like receptor signaling defects, over activation of neutrophil subsets, decreased regulatory T cell frequency due to T cell tolerance defects, excessive type I interferon downstream activation, and autoantibody production by pathogenic B cells, to name a few (Crow, 2014; Zharkova *et al*., 2017). Therefore, one should expect that study of a single or a couple immune cell subsets in isolation would not fully depict the immunopathology. Recent publications have supported the multi-immune component etiology of the disease, and in fact, different disease phenotypes may be correlated with specific immune profiles that encompass the integrated immunological picture (Nehar-Belaid *et al*., 2020). Therefore, our work analyzing multiple immune cell subsets and their downstream cytokine production, as a composite integrated approach, most accurately addresses the basic science and clinical questions.

Each of the datasets described in the “Motivating Data” section were generated via Fluidigm mass cytometer instruments. Resultant FCS files were bead normalized to account for instrument calibration. The final cell-type abundance proportion matrix after the data filtering steps (debarcoding, bead normalization, batch adjustment/normalization) had 28 rows and 12 columns, where each row corresponded to an observation *P*^(*i*)^ *∈* S^12^ and each column to a cell type. These different cell types were CD4+ T-cells, CD8+ T-cells, CD27-hi B-cells, CD27-lo B-cells, Basophils, CD14-hi monocytess, CD16-hi monocytes, CD56-hi NK-cells, CD56-mid NK-cells, cDCs, pDCs and Other cells. The predictor of interest was the disease status (SLE or healthy control), and a potential covariate was the stimulation condition (T0 or T6). Appyling the CODAK test on the full data set with 10^6^ permutations resulted in a distance correlation 0.6966 and a p-value *<* 10^−6^. When adjusted for the covariate stimulation condition (using CODAK-sk), the distance correlation was 0.7974 and the p-value *<* 10^−6^. However, one needs to be cautious due to the fact that the study contains 2 repeated measures on each individual subject which leads to the violation of the independence assumption. Separate testing for conditions T0 and T6 still resulted in statistically significant results (*P* = 0.0006 for both T0 and T6).

We also obtained the *dcor*_*LOO*_ statistic for every cell type. The results are shown in Figure 7. The top 3 contributors for both conditions were CD27-lo B-cells, conventional dendritic cells (cDCs), and CD16-hi monocytes. These results are consistent with previously published literature. Pathogenic B-cells have long been implicated in the pathogenesis of SLE, including expansion of CD21-lo B cells, which are also most often CD27-lo (Dörner *et al*., 2011). Monocytes and dendritic cells constitute key elements of a pro-inflammatory innate immune response and they are significantly influenced by the circulating cytokine environment. In the study group shown, patients demonstrated significantly increased pro-inflammatory serum cytokines (data not shown), which likely accounts for the expanded CD16-hi monocytes and cDCs subpopulations (Steinbach *et al*., 2000).

**Figure 7:**
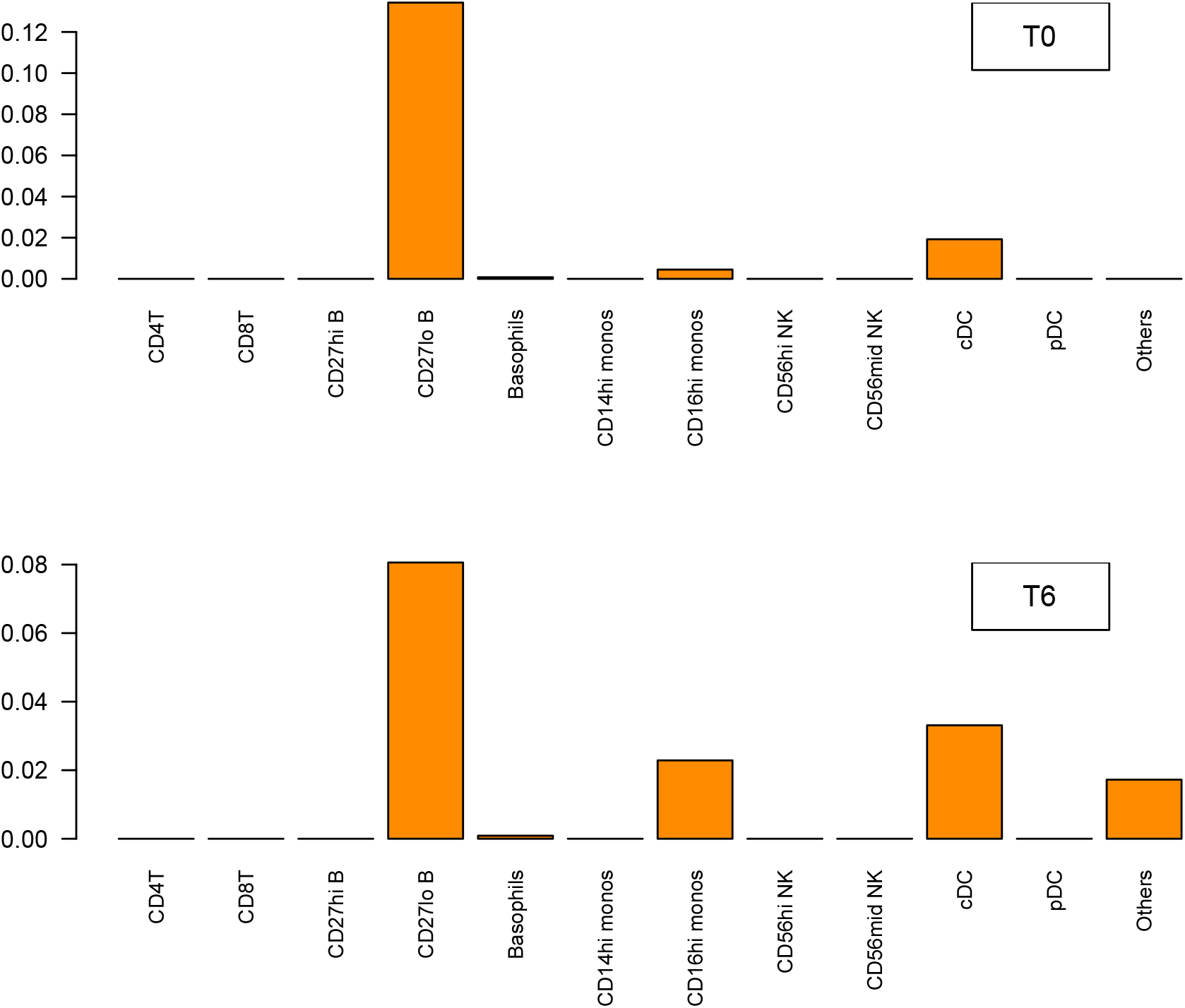
The values of the *dcor*_*LOO*_ statistic for testing the difference in cell type abundance when comparing SLE vs healthy controls at T0 and T6.

Please see Supplementary Materials (section S2) for a second data analysis example containing a continuous predictor (study 2 from motivating data).

## 4 Discussion

We proposed an appropriate kernel to quantify similarities in the cell-type abundance profiles for mass cytometry data using the AD. The proposed kernel enjoys the desirable properties of the AD, and also performs well for both simulated and real data sets. Unlike some exisiting methods, our framework CODAK can also adjust for covariates, both categorical and continuous.

One common issue for testing cell-type abundance for CyTOF is that the number of cell types can often be large and the sample sizes are often small. Many statistical tests, e.g. the tests for the GLMM, are asymptotic tests that do not perform well for such small sample scenarios. In particular, these tests are often anticonservative while resampling versions of these tests can be computationally intensive. We have shown that other tests not specifically requiring large samples in theory can also be anti-conservative. Our simulation studies demonstrate that CODAK has much superior type-I error control and competitive power when compared to the state of the art methods. CODAK is also nonparametric in nature, and therefore expected to be more robust compared to existing parametric models against model violations such as violation of distributional assumptions. In one example we even show that the CODAK-sk method is somewhat robust against violation of the independence assumption and can be used, with caution, for repeated measures data if the repeated measures are expected to be only mildly correlated.

Another important feature of CODAK is that it tests for the global null hypothesis of association of the predictors with the full compositional profile of the cell types, and therefore does not suffer from the burden of multiple testing. Such gain is most significant when a number of cell types are associated with the predictor variable. Figure 4 provides an insight into when this approach is most avdantageous compared to univariate testing using GLMM or diffcyt-voom. CODAK is also able to test association for continuous predictors while some other methods (e.g. diffcyt) do not provide an easy way to do so.

In this article, we have shown the ability of kernel machines to have high power for testing for compositional associations with clinical outcomes, which mirrors its success in other settings (e.g., Wu *et al.* (2011); Rudra *et al.* (2018)). Nevertheless, because the methodology operates at the level of samples and not individual features, one criticism is its interpretability. As suggested in Section 2.3., one can follow up using existing component-level tests and use gatekeeper type methods to adjust for multiple comparisons. We also suggested two kernel based approaches to rank the individual features in order of their contribution to the overall association. Another finding is that the proposed method is powerful when there are small differences in the abundance of several cell types. By contrast, if only a small subset of cell types show a difference, the simulations show that our method loses power. However, in immunological disease processes, most pathology affects multiple immune cell subsets and different downstream functional effects (Waugh *et al*., 2019; Galbraith *et al*., 2021). Finally, our methodology cannot handle zero proportions. While there exist techniques from the compositional data literature to handle this (replacing zeros by a small number, for example), they are beyond the scope of the current manuscript.

## 5 Conclusion

We have developed a statistical framework based on kernel distance covariance to test association between compositional profiles of cell type abundance with important predictors for mass cytometry data. Our framework can scale up well for high-dimensions and performs well even in small samples. We also proposed methods for covariate adjustment as well as follow-up methods for finding the top cell types contributing to the association. Using extensive simulation studies, the method has been shown to perform well compared to the existing methods under many scenarios. We also demonstrated the performance of the method in real mass cytometry data sets. The approach has further potential to find application for more complex applications such as immunogenomics for multidimensional predictors. With rising applications of CyTOF, our framework provides an important contribution towards the analysis of high-dimensional cell-type abundance data.

