## Supplementary Materials for "Compositional Data Analysis using Kernels in Mass Cytometry Data"

#### S1 Additional Simulations

##### S1.1 Simulation details

We conducted the following simulation studies besides the ones reported in the main manuscript.

S(i) Continuous predictor: This is identical to scenario (i) in the main manuscript (Figure 3) except that the predictor variable is a continuous variable simulated from a $U(0, 1)$ distribution. For comparability, the effect sizes in cases (a)-(d) are simulated such that the mean and variance of $\beta X$ remains same as the binary predictor case, where $\beta$ is the $q$-dimensional vector of effect sizes.

S(ii) Binary predictor with larger random effects variance: We also wanted to explore the effect of a larger random effect variance. This simulation is identical to the to scenario (i) in the main manuscript (Figure 3) except that the standard deviation of the random effects term is now assumed to be 0.5.

S(iii) Continuous predictor, binary covariate: This is similar to case (ii) in the main manuscript (Figure 5) except that the predictor is now considered to be continuous like S(i) above.

##### S1.2 Results of the additional simulations

The comparisons of the size and power for the methods under consideration for simulations S(i)-S(iii) are shown in Figures S1, S2, S3, respectively. The diffcyt methods were not available for the continuous predictor cases. The findings are similar to the simulations in the main manuscript. The advantage of CODAK over the other two methods slightly diminished when a larger random effects variance was used.

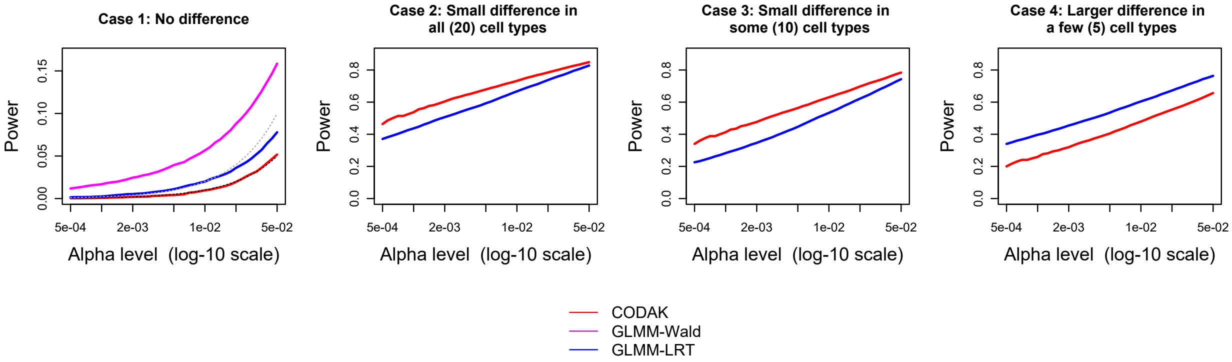


Figure S1: Comparison of statistical power for continuous predictor. The black dashed line in the first plot shows the nominal level $\alpha$ and the grey dashed line shows two times $\alpha$. Only the methods with reasonable control of type-I error are shown in the other 3 plots.

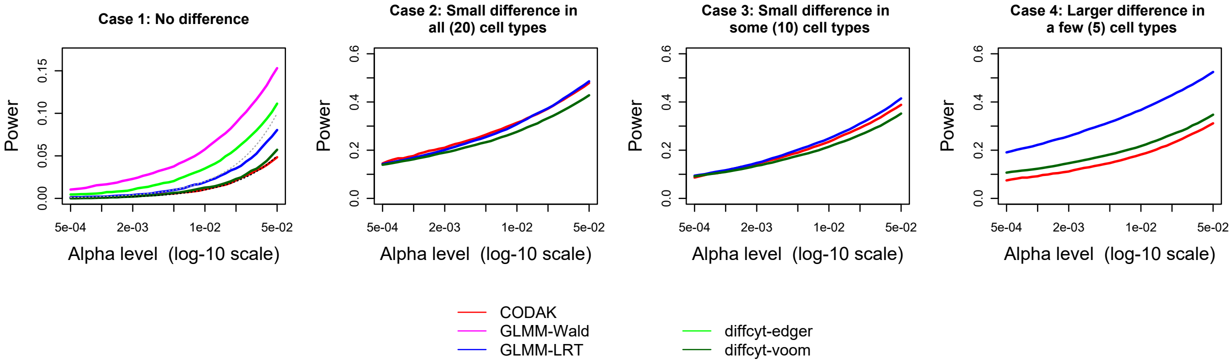


Figure S2: Comparison of statistical power for larger random effects variance (binary predictor). The black dashed line in the first plot shows the nominal level $\alpha$ and the grey dashed line shows $2\alpha$. Only the methods with reasonable control of type-I error are shown in the other 3 plots.

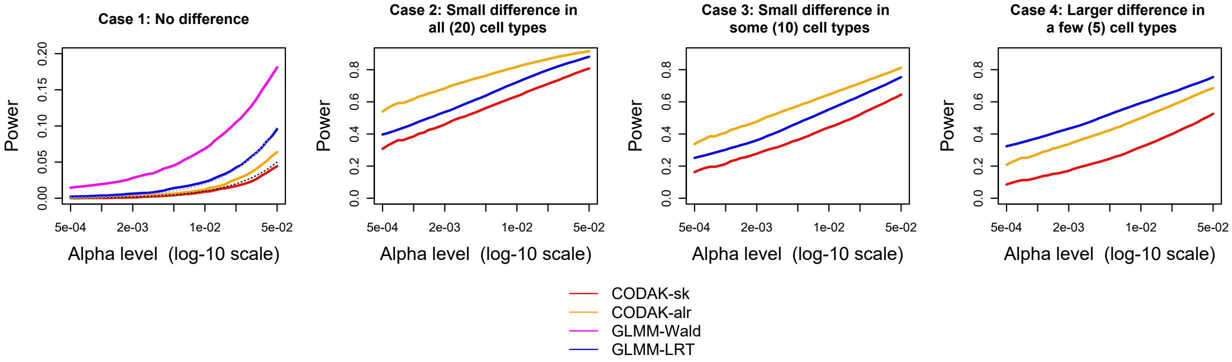


Figure S3: Comparison of statistical power for continuous predictor, categorical covariate. The black dashed line in the first plot shows the nominal level $\alpha$ and the grey dashed line shows $2\alpha$. Only the methods with reasonable control of type-I error are shown in the other 3 plots.

| | |
|---|---|
| CD4 + T cells | |
| 1 | CD4 + Naïve T cells |
| 2 | CD4 + Central memory T cells |
| 3 | CD4 + TEMRA |
| 4 | CD4 + Effector memory T cells |
| CD8 + T cells | |
| 1 | CD8 + Naïve T cells |
| 2 | CD8 + Central memory T cells |
| 3 | CD8 + TEMRA |
| 4 | CD8 + Effector memory T cells |
| Non T Non B cells | |
| 1 | CD45RA-CD56- |
| 2 | CD45RA-CD56+ NK |
| 3 | CD11c + Myeloids |
| 4 | HLADR + CD11c- Monocytes |
| 5 | CD56dim CD16 + NK |
| 6 | CD56dim NK |
| 7 | CD56hi NK |
| 8 | CD14hi Monocytes |
| 9 | CD16hi Monocytes |
| 10 | CD14- CD16- |
| 11 | pDCs |
| 12 | mDCs |
| CD19 + B cells | |
| 1 | CD27hi B cells |
| 2 | CD27lo B cells |
| 3 | IgM + IgD + Pre-switched |
| 4 | IgM + IgD - IgM Memory |
| 5 | IgM - IgD + c-delta class switched |
| 6 | IgM - IgD - Switched Memory |
| 7 | IgM + IgD + Mature Naive |
| 8 | IgM - IgD + Immature Naive |
| 9 | IgM - IgD + Anergic B cells |
| 10 | IgM - IgD – Atypical memory |

Figure S4: A complete list of cell type names for study 2 (corresponds to Figure 1 in the main manuscript)

### S2 A second data analysis example

This analysis is based on data collected from an ongoing study conducted by the authors (study 2). Peripheral whole blood sample was collected from SLE patients and age and sex-matched controls. Peripheral blood samples were fixed (and red blood cells lysed) either immediately after collection (T0); or after incubation at 37C with a protein transport inhibitor cocktail (T6). Lysed/fixed cells were stored at $-80°C$, and were thawed on the day of barcoding and staining. To decrease technical variability, palladium isotopes were used in different combinations for mass tag barcoding of separate samples, pooled in sets of 20, surface stained in a single tube with a metal-labeled antibody panel. Barcoding methodology was adapted from Zunder *et al.* (2015). Protocols for intracellular cytokine staining (ICS) assays were adapted from previous studies in O'Gorman *et al.* (2017). Individual samples were barcoded to reduce technical variability, and data were also batch adjusted (Schuyler *et al.*, 2019). Clinical data including clinical laboratory parameters (complete blood cell counts, autoantibodies, chemistry panels, etc), clinical disease manifestations identified on physical exam, imaging studies, and disease severity scores (SLE disease activity index, SLEDAI) were recorded from every participant and every study timepoint (Bombardier *et al.*, 1992; Touma *et al.*, 2011). Nephritis diagnosis, to be used as a covariate, was based on renal biopsy pathology evaluation (Schwartz *et al.*, 2014).

Although questions similar to study 1 are relevant for this study too (e.g. SLE vs healthy control comparison), in this paper, for the purpose of demonstration of the application of CODAK for a continuous predictor, we focused on the association between the cell-type abundance and the SLEDAI score, a continuous predictor. Since we are interested in the disease activity (SLEDAI score), we only used the data from the SLE patients at T0. Application of CODAK on this data resulted in a p-value = 0.2794 showing that there is no significant association of the cell type compositions with the SLEDAI score. We also conducted analysis while adjusting for the effect of the binary covariate whether an SLE patient had nephritis or not. CODAK-sk and CODAK-alr both resulted in statistically non-significant outcomes (p-value = 0.3914 and 0.4298, respectively).

### S3 Proof of the result in Equation 4

To show:

$$\underset{c}{\operatorname{argmin}}\ d(P^{(i)}_{-c}, P^{(j)}_{-c}) = \underset{c}{\operatorname{argmax}} \left| \ln\left[\frac{P_{ic}}{P_{jc}}\right] - \ln\left[\frac{g(P^{(i)})}{g(P^{(j)})}\right] \right|. \tag{S1}$$

*Proof.* It is easy to see that the Aitchison Distance (AD) between the two compositions $P^{(i)}$ and $P^{(j)}$ can be written as:

$$d(P^{(i)}, P^{(j)}) = \sum_{r=1}^{q} (V_r - \bar{V})^2,$$

where $V_r = \ln\left[\frac{P_{ir}}{P_{jr}}\right]$, by realizing that

$$\bar{V} = \frac{1}{q}\sum_{r=1}^{q} \ln\left[\frac{P_{ir}}{P_{jr}}\right] = \ln\left[\frac{g(P^{(i)})}{g(P^{(j)})}\right].$$

In the light of this, Equation S1 becomes

$$\underset{c}{\operatorname{argmin}} \sum_{r\neq c}(V_r - \bar{V}_{-c})^2 = \underset{c}{\operatorname{argmax}} \left|V_c - \bar{V}\right|, \tag{S2}$$

where $\bar{V}_{-c} = \frac{1}{q-1}\sum_{r\neq c} V_r$.

Now,

$$\begin{aligned}
\sum_{r\neq c}(V_r - \bar{V})^2 &= \sum_{r\neq c}(V_r - \bar{V}_{-c})^2 + q(\bar{V} - \bar{V}_{-c})^2 \\
\Rightarrow \sum_{r\neq c}(V_r - \bar{V}_{-c})^2 &= \sum_{r\neq c}(V_r - \bar{V})^2 - q(\bar{V} - \bar{V}_{-c})^2 \\
&= \sum_{r=1}^{q}(V_r - \bar{V})^2 - (V_c - \bar{V})^2 - q(\bar{V} - \bar{V}_{-c})^2 \\
&= \sum_{r=1}^{q}(V_r - \bar{V})^2 - (V_c - \bar{V})^2 - \frac{q}{(q-1)^2}(V_c - \bar{V})^2
\end{aligned}$$

Note that the two rightmost terms in this last equation are both increasing functions of $|V_c - \bar{V}|$. This completes the proof. □
